# MOAL: Multi-Omic Analysis at Lab. A simplified methodology workflow to make reproducible omic bioanalysis

**DOI:** 10.1101/2023.10.17.562686

**Authors:** Florent Dumont, Emy Ponsardin, Guillaume Bernadat, Sylvia Cohen-Kaminsky

## Abstract

Omic data research projects are still developing today, but analysis suffers from not having a standardized workflow for biostatistics and functional integration steps that we gather here as bioanalysis. To make bioanalysis more reproducible and accessible, we created MOAL (Multi-Omic Analysis at Lab), an R package including a omic() function that automates most classical tasks. We reproduced in one go the results of a published dataset starting from unfiltered available data confirming the capabilities to automate the production of omic bioanalysis using MOAL.

## Context

### Omic bioanalysis features

Numerous efficient bioinformatics tools are available for preprocessing and generating read-outs measured matrix. Typically, the following step is normalization, which is, similarly, assisted by numerous mathematical methodologies available in programming or graphical ways. At this step, we usually must deal with a numerical normalized matrix unballasted from technical device noise and ready to make downstream analysis. We define as « bioanalysis » final omic analysis steps leading to biological interpretation scaffolding by infographics integrated into scientific literature data.

Bioanalysis could be summed up by the biostatistics step, including descriptive and differential analysis followed by functional (or enrichment) analysis: biostatistics aims at extracting relevant subsets from the global dataset, and functional analysis seeks to help interpretation by integrating these subsets into biological and medical knowledge. To facilitate integration, knowledge is summarized and organized in gene sets. We can define gene sets as a unity of knowledge that can come from: 1) ontologies that link biological functions in a hierarchy, 2) canonical pathways that link genes (or their products) supported by robust, reliable, and reproducible experiments across literature, or 3) simple validated list of genes associated by statistical correlation with a transcription factor, a miRNA, a long coding RNA, a clinical trait, a phenotype, an immunological or an oncological pattern.

Bioanalysis suffers from not having a simple, standardized, and automatized workflow for reproducible research at the laboratory. In this context, we developed MOAL (Multi Omic Analysis at Lab) a R package including an easy-to-use omic function in response to this lack.

### Source of variability

The aim of an experimentation and its analysis is to find biological insights by reducing technical noises to better estimate the individual variability coming from the biodiversity of a population. Statistics quantify this variability in an undirected way by using a subset of the population, thus, the p-value computed starting from the sample size and the observed fold-change can only say if the sample size is large enough to correctly represent the population variability to consider that a difference exists. Consequently, observatories use statistical methods as a standard and produce descriptors incorporating a quantified risk, which enables them to conclude regarding the presence or absence of a phenomenon at a biological level, and possibly compare their conclusions with other results obtained under close conditions.

Each step of a complete omic analysis workflow presents its source of variability. Fig1. Illustrate and sum up the source of variability that one could meet at each step of an omic project. First, the experimental step asks for an appropriate design of the number of treatment factors, the number of levels by factor, and the number of samples by level. The more sample we have by groups, the more we can avoid confusing variables with the help of contingency tables to see sample distribution across all factors’ level combinations. It is also important to know if samples are linked by pairing (i.e., two measurements on the same subject) or by nested factors (i.e., when the design requires combination levels between two factors with no subjects). The last case to consider is about the interaction between two factors that could be included in the model, enabling to distinguish a synergy from a cumulative effect. All these cases cover a major part of experimental design and can be modeled by analysis of variance. The next step is to pass sample extracts in a high-throughput device, which is already known to have some variability across instrument models and host platforms. At this step, formatted files including device parameters, metadata, metrics, raw data signals, and possibly systemic biases are available. Because of the diversity in omic technologies, bioinformatic preprocessing methodologies and tools are varied and numerous. For example, bioinformatic practices are very different in getting the numerical matrix containing raw signals when dealing with mass spectrometry .raw files or .fastq sequencing files. Preprocessing also includes raw data normalization procedures, which aim at increasing signal/noise ratio and make data more comparable by deleting systemic biases from samples or variables. At this step, three tables are generally accessible for omic bioanalysis: the normalized data file and the sample information file including experimental, phenotypic, clinical and design factors for biostatistics. The third table is an annotation file including at least a Symbol column and an identifier column matching with variables.

All methodologies described above have their contribution to variability in an omic project. Here, we automate the two final steps called bioanalysis corresponding to biostatistics and functional integration procedures. The aim of standardizing bioanalysis is to use biologically interpreted results to have quality indicators that will help understanding and standardizing upstream experimental biases and ultimately emancipate from variables with little latitude such as device types, parameters and plateform practices. Here, we chose an unfiltered published data set [1] to test our omic workflow and see its capacities to generate reproducible and interpretable biological results in one go on a laptop.

## Material and methods

First, the workflow used descriptive statistical tools for quality controls: histogram and boxplot for distribution across samples, PCA and HCA for outliers and relevant factors, Fratio barplot for ANOVA model.

As mentioned above, we apply a classical analysis of variance for differential analysis of data. ANOVA belongs to the linear model test family and is a simple and useful way to adjust for most experimental designs. It allows keeping all samples in the same statistical model, which holds statistical power for residual variance estimation and enables comparing biological and technical factor variability from one to another. It also increases fold-change contrast between comparisons because, in the ANOVA context, means are computed as a linear combination using the variance estimation across treatment groups in accordance with the experimental design. Furthermore, we implemented the remove batch effect function from the limma package [2], thus allowing deletion of technical bias that was correctly balanced across factor combinations during sample preparation and guaranteed a better feature selection and clustering for downstream analysis.

For annotation tasks, symbols are automatically re-annotated using synonym checking to avoid information loss. We also integrated the NBCI orthologs gene database to open functional enrichment analysis for species that have identified ortholog genes in human. Consequently, we created additional annotation packages to update all bibliographic data simultaneously, improve results reproducibility, and decrease computing time. Annotation databases include NCBI gene [3] for 11 species,^1^ NCBI orthologs [3], Ensembl [4], and Uniprot [5]. Gene sets databases include MSigDB [6,7] and the Gene ontology [8]. StringDB [9] is used for interaction network.

Dependency packages providing notably language extensions, parallel processing, analytical technique-specific, clustering, functional analysis and graphing functions have been used [2,10-40].

Fig.2 shows all biostatistics and functional analysis tasks realized by the omic() function workflow.

## Results

### Application example: re-analysis of the published dataset GSE65055

As a positive control, we reproduced the enrichment analysis of transcriptomic analysis carried out on aneuploidy samples for trisomy 13, 18, 21 and the associated control samples [1] (https://pubmed.ncbi.nlm.nih.gov/27283765/).

The study infers that “overall genome expression profiles highlighted changes in the expression of a subset of genes in trisomic chromosomes, while the majority of transcriptional changes concerned genes located on euploid chromosomes.” [1] (Fig. 2)

**Fig. 1:**
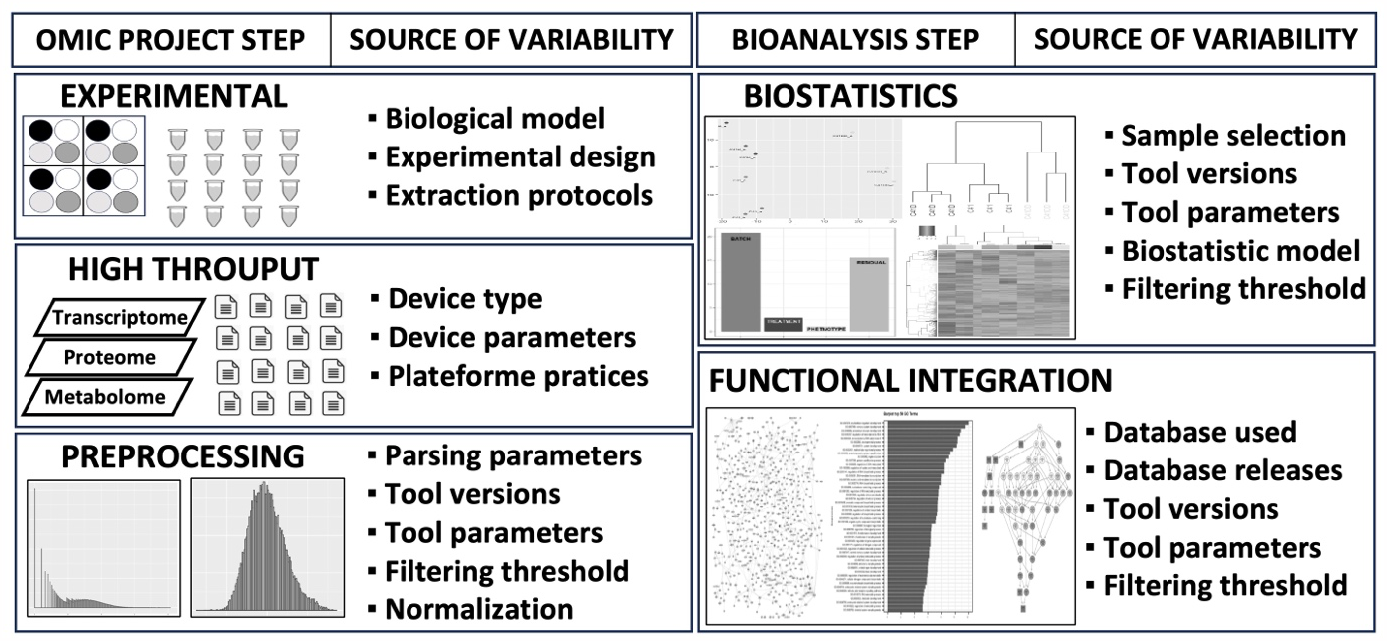
Sources of variabilty for each omic project step are coming from protocols, plateform pratices as well as all computational methodologies used to produce complete interpretable biological results.

**Fig. 2:**
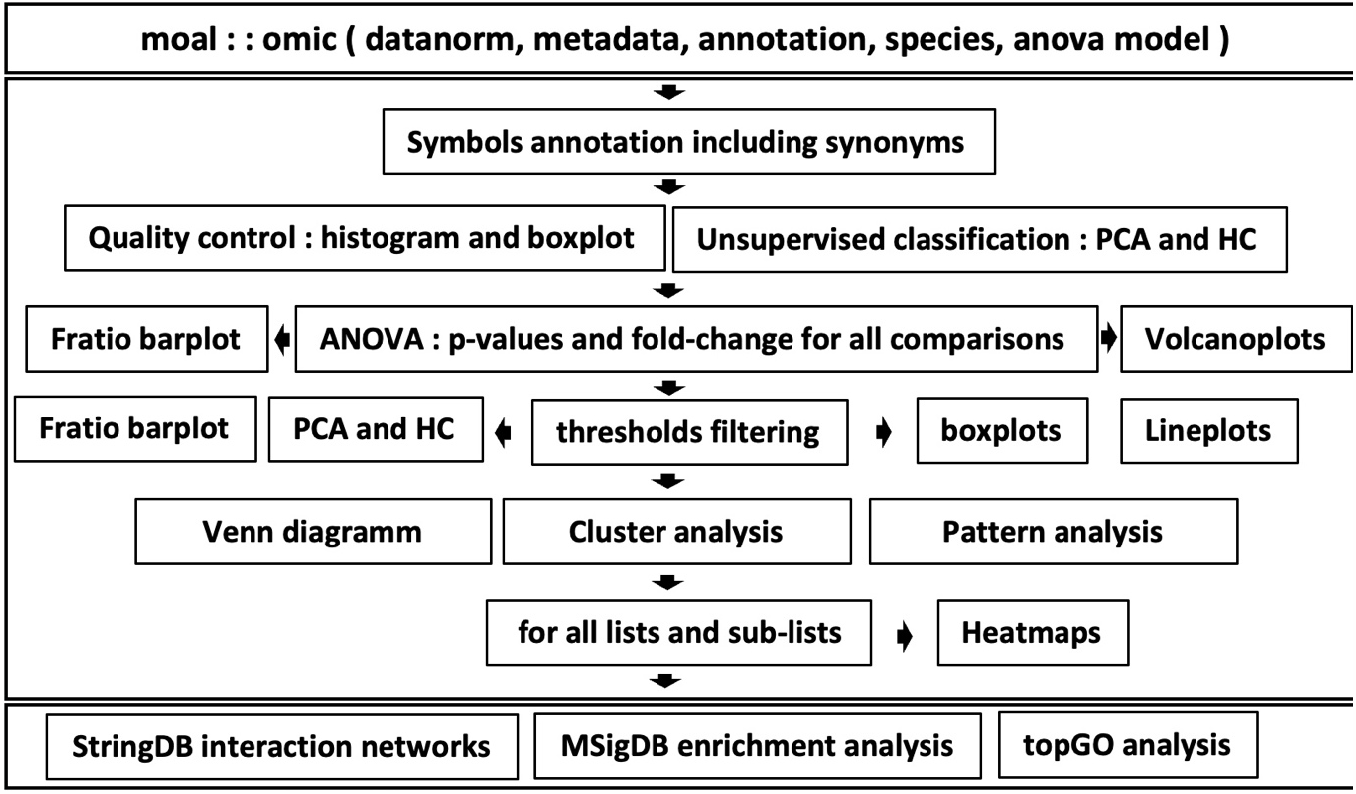
Summary of bioanalysis workflow processed by omic() function.

**Fig. 3.**
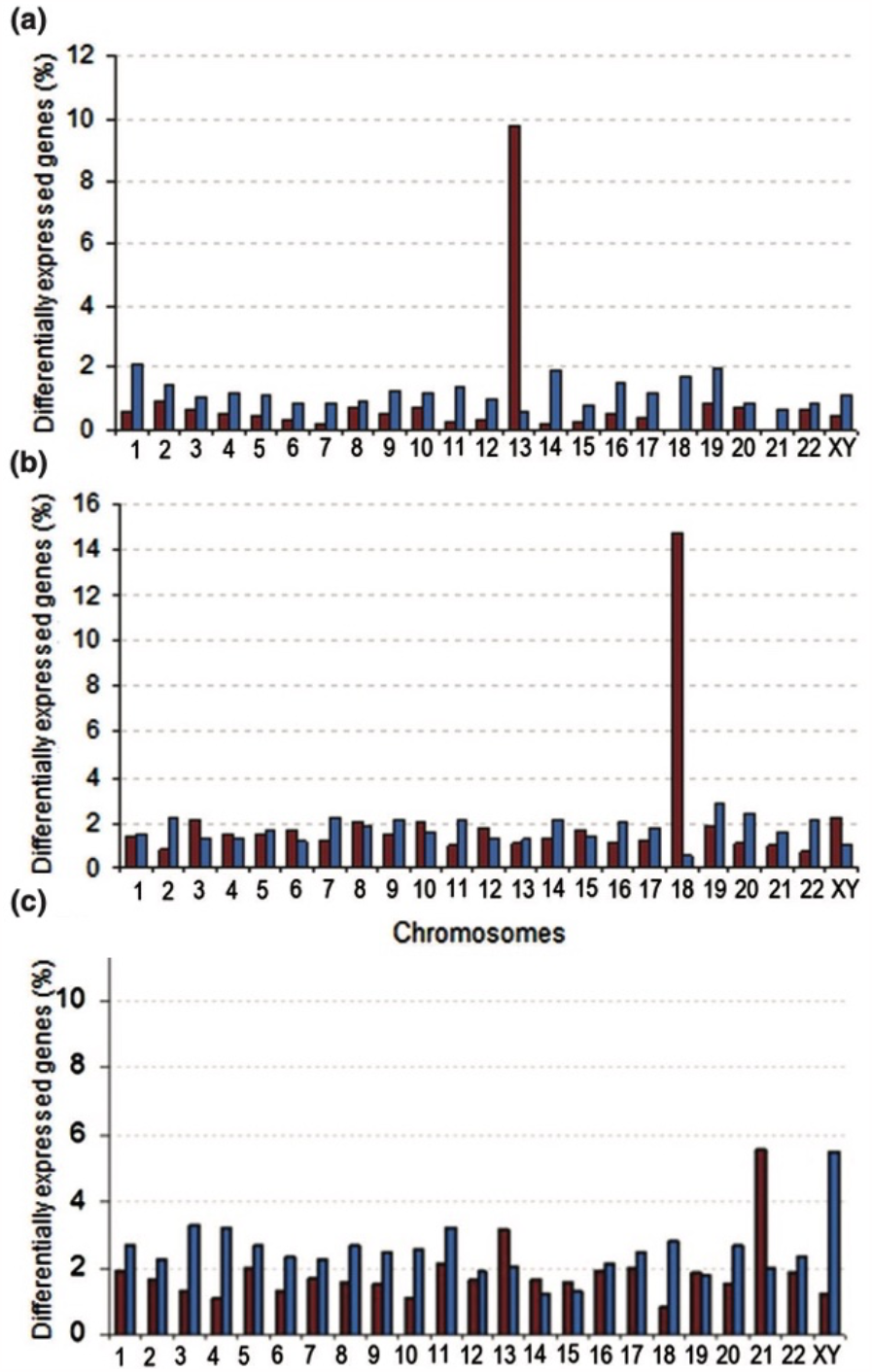
Enrichment analysis results presented in the publication Hervé et al [1]. Percentages of differentially expressed genes sets for each chromosome is indicated on the y-axis and show an overrepresentation of dysregulated genes for trisomic chromosomes: (a) for trisomy 13, (b) for trisomy 18 and (c) for trisomy 21).

### Quality controls

“We therefore chose only to compare expression profiles produced from the same type of sample to avoid any confusion between aneuploidy effect and cell-type specific differences.” “Lastly, we compared five trisomy 21 chorionic villus samples with four euploid chorionic villus samples.” [1]

As explained in the publication above, one outlier sample was removed from chorionic tissue (GSM1586484), and we divided the experimental design into two ANOVA models for each tissue type because of the heterogeneity. Fig. 4. shows the replication of these two observations using the data processing matrix available from GEO.

**Fig. 4.**
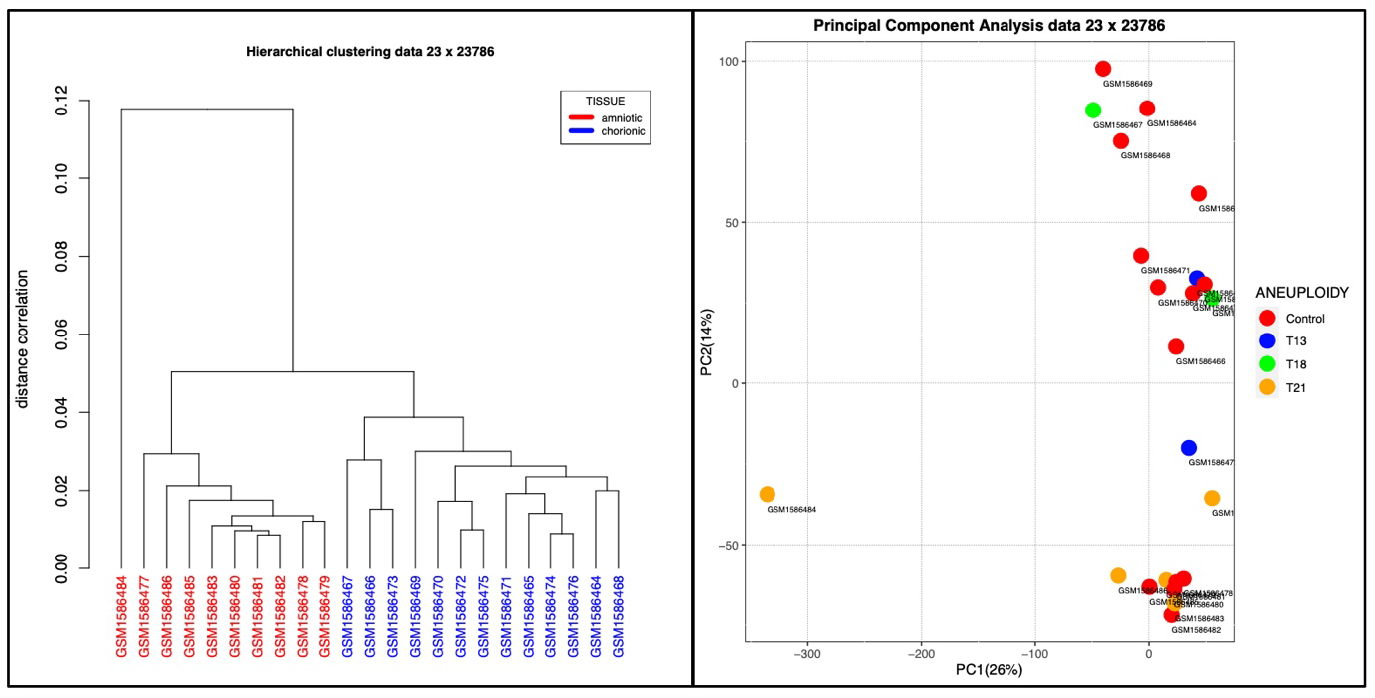
Quality controls of the re-analyzed data set showing one outlier sample GSM1586484 removed for downstream analysis and main effect of tissue represented by the amniotic sample cluster in red and chorionic cluster in blue.

### Differential analysis

As illustrated in Fig. 5., in order to reproduce the differential analysis, we started from the same unfiltered matrix and chose to apply a unique 2-ways ANOVA model including ANEUPLOIDY and TISSUE factors instead of the two separated analyses as described above.

**Fig. 5.**
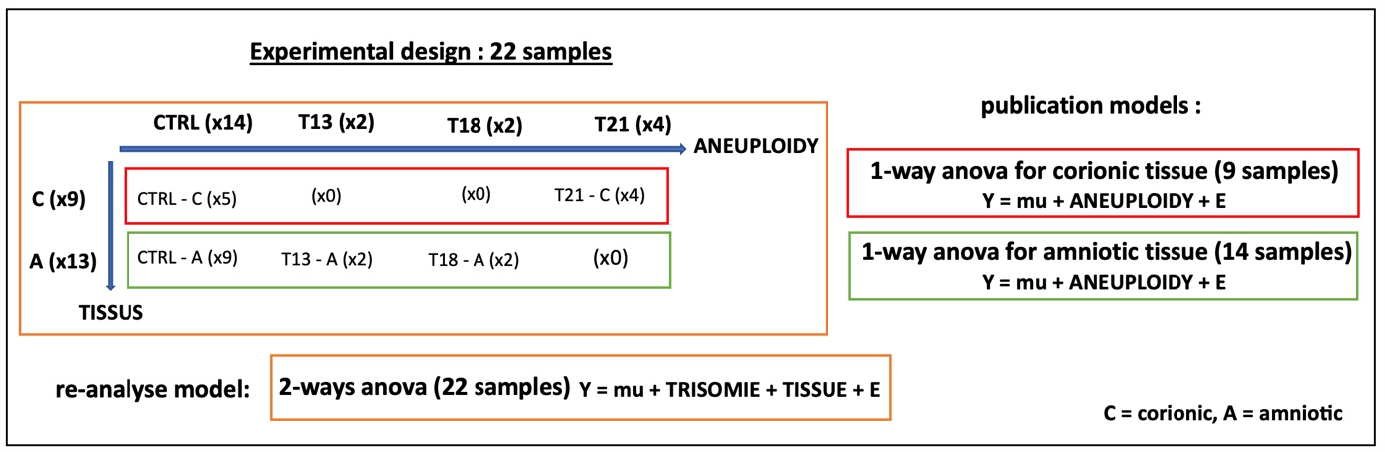
Experimental design and ANOVA models applied on re-analyzed data.

### Enrichment analysis

As shown in Fig.6, we reproduced enrichment results starting from unfiltered matrix data using the same thresholds (p-value < 0.05 and fold-change > 1.5). Here, using MSigDB positional gene set collection corresponding to human chromosome cytogenetic bands, we found the best enrichments and p-values for aneuploid chromosomes corresponding to the up-dysregulated list of genes.

**Fig. 6.**
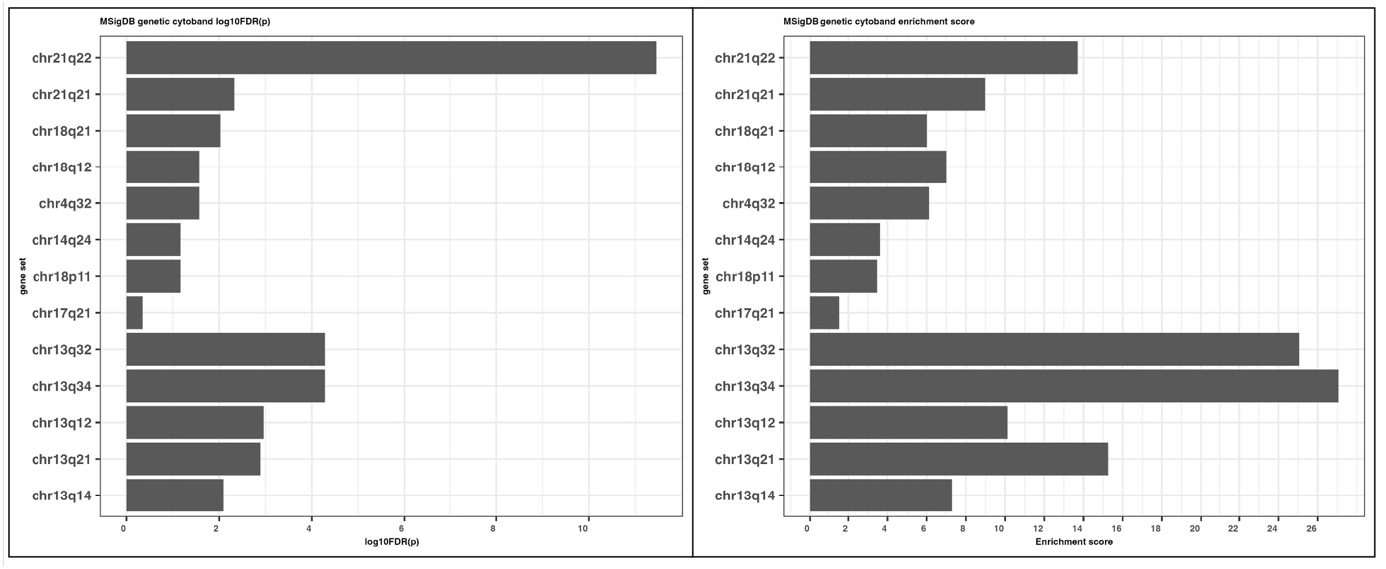
Enrichment analysis result obtain for re-analyzed data showing significative gene sets belonging to trisomic chromosomes as observed in the publication.

We observed that few other gene set cytobands give significative enrichment scores for euploid chromosomes. More precisely, starting from 299 gene sets in the collection, we obtained 10 significative gene sets for trisomic chromosomes and 3 for others (chr4q32, chr14q24, chr17q21). These results validate the re-analysis of the dataset and workflow capabilities to produce relevant biological interpretations starting from unfiltered data.

## Discussion

The R package we have developed aimed at showing that it was possible to automate and integrate classical biostatistical and functional omic analysis available for a dozen of current biological models, as well as illustrating the advantages of shortening and alleviating the process leading up to biological interpretation.

Starting simply from an unfiltered normalized data table and its experimental design, one easy-to-use function enabled generating a complete (unsupervised and supervised) analysis. Results contain PCA and HCA plots, which allow observation of the distribution of samples across treatment groups without *a priori*, as well as ANOVA F ratio barplot, which helps evaluating quality of model adjustment with the experimental design.

By interrogating classical biological and medical knowledge databases based on relevant features at several threshold levels and displaying results using simple infographics, we hope to have contributed some form of one-stop computer-aided biological interpretation. By leveraging ortholog information for extra species, we are also trying to provide tools that facilitate reasoning beyond the classic rodent and human models.

Output is organized in a directory with an unchanged structure between runs (favoring open, easy-to-read and reusable file formats), while attempting to capture input parameters and database versions, as well as recording graphics and tables produced to facilitate focus on significant information. One can, therefore, change a unique parameter and quickly determine its influence on results from the biological standpoint.

Typically, outliers can be detected/excluded, and unexpected batch factor can be added to the ANOVA model, before the analysis is re-run at little cost, which facilitates comparison of results based on biological interpretation.

Moreover, all discriminant patterns generated at several threshold levels can be easily compared simultaneously at the functional level, which allows one to be critical about the influence of statical thresholding on biological interpretation.

(Parts of) our package could also prove helpful to control and prepare data for other workflows. For example, when removing a batch effect (or matching factor) according to the experimental design is required, the unballasted matrix ready for downstream analysis can be retrieved from the results directory.

Regarding annotation and species, symbols are automatically annotated using synonym checking, and functional analysis can be carried out for all species that have identified ortholog genes in the human NCBI gene database.

## Conclusion

In the context of multi-omic data analysis, we found no such tools gathering current biostatistical and functional analysis tasks in the same workflow to produce biologically enriched results locally in one go. We attempted to fill this gap with a R package containing a single, easy-to-use function requiring only an unfiltered normalized data table and the corresponding experimental design. We expect this tool to create a higher level of experimentation regarding the data processing phase, thereby facilitating and improving the optimisation of related parameters and contributing to maximize scientific yield of input data.

## Install

For users interested in using the software immediately: binary packages and instructions for installing are currently available from: https://www.ipsit.universite-paris-saclay.fr/?-bioinfo-(a https://fdumbioinfo.r-universe.dev subdomain is in progress).

For developers/biostatisticians: source files and instructions for building (update annotations) and installing packages are available from Zenodo: https://doi.org/10.5281/zenodo.10013866.

## Abbreviations

PCA: Principal Component Analysis,
AHC: Ascendant Hierarchical Clustering,
ANOVA: analysis of variance

## Acknowledgements

We thank Claudine Deloménie for helpful discussions and testing the MOAL package, Paul Savescu for testing MOAL and developing future features for helping results interpretation, and Françoise Cormier for her help in testing MOAL and interpretating biological results, all from the IPSIT **(Ingénierie et Plateformes au Service de l’Innovation Thérapeutique)** team. IPSIT belongs to Université-Paris-Saclay. IPSIT is a UMS (Unité Mixte de Service US31 Inserm, UAR3679 CNRS), which brings together technological platforms. IPSIT intends to be firmly at the interface of chemistry, biology, and medicine, establishing the link between the pathological target and the drug. Thanks to Laura Samrani for help testing topGO biological results analysis from UMR-S 996 at Paris-Saclay University. The Migale Bioinformatics facility, GaBi @BRIDGe (Université Paris-Saclay, INRAE, 78350, Jouy-en-Josas, France) platform and UPSay IT department are gratefully acknowledged for computational resources, technical support and/or statistical advice.

## Footnotes

1 *Bos taurus, Caenorhabditis elegans, Danio rerio, Drosophila melanogaster, Gallus gallus, Homo sapiens, Mus musculus, Pan troglodytes, Rattus norvegicus, Sus scrofa, Xenopus tropicalis*.

